# A haplotype resolved chromosome-scale assembly of North American wild apple *Malus fusca* and comparative genomics of the fire blight *Mfu10* locus. Genome of the Pacific Crabapple *Malus fusca*

**DOI:** 10.1101/2023.03.22.533842

**Authors:** Ben N. Mansfeld, Alan Yocca, Shujun Ou, Alex Harkess, Erik Burchard, Benjamin Gutierrez, Steve van Nocker, Christopher Gottschalk

## Abstract

The Pacific crabapple (*Malus fusca*) is a wild relative of the commercial apple (*Malus* × *domestica*). With a range extending from Alaska to Northern California, *M. fusca* is extremely hardy and disease resistant. The species represents an untapped genetic resource for development of new apple cultivars with enhanced stress resistance. However, gene discovery and utilization of *M. fusca* has been hampered by the lack of genomic resources. Here, we present a high-quality, haplotype-resolved, chromosome-scale genome assembly and annotation for *M. fusca*. The genome was assembled using high-fidelity long-reads and scaffolded using genetic maps and high-throughput chromatin conformation capture sequencing, resulting in one of the most contiguous apple genomes to date. We annotated the genome using public transcriptomic data from the same species taken from diverse plant structures and developmental stages. Using this assembly, we explored haplotypic structural variation within the genome of *M. fusca,* identifying thousands of large variants. We further showed high sequence co-linearity with other domesticated and wild *Malus* species. Finally, we resolve a known quantitative trait locus associated with resistance to fire blight (*Erwinia amylovora*). Insights gained from the assembly of a reference-quality genome of this hardy wild apple relative will be invaluable as a tool to facilitate DNA-informed introgression breeding.

## Introduction

The Pacific crabapple (*Malus fusca*), one of four native North American species, is found in the Pacific Northwest ranging from Alaska and British Columbia to California (USDA Agricultural Research Service, 2015). These hardy trees routinely grow in conditions in which the vast majority of cultivated apple (*Malus* × *domestica* Borkh.) cultivars cannot survive and reproduce in; *M. fusca* can withstand winters of −46°C or colder (Fiala 1994), can grow on beach heads, in sandy soils, exposed to brackish water and in waterlogged conditions (USDA Agricultural Research Service, 2015; Volk, 2019). Moreover, *M. fusca* has been found to be resistant to fire blight (*Erwinia amylovora*), a devastating disease that is endemic to North America, but now found worldwide (Dougherty et al., 2021; Emeriewen et al., 2014; Bonn & van der Zwet, 2001; Vanneste, 2001).

Many environmental conditions, including those derived by human caused climate change, increasingly burden apple production (Volk et al., 2015a). For example, abiotic stress such as water logging and spring frosts during bloom can lead to reduced yields, disrupted growth patterns, and in extreme situations, loss of trees (Atkinson et al., 2000; Bhusal et al., 2019; Dalhaus et al., 2020; Gottschalk and van Nocker 2013, Schrader et al., 2001; Torres et al., 2013, 2016, Way et al., 1991). Furthermore, in addition to abiotic stresses, apple (*Malus* × *domestica*) suffers from other devastating diseases apart from fire blight, such as apple scab, bitter pit, and cedar apple rust, as well as other pests (*e.g.*, codling moth). These can additionally reduce yields or the market value of fruit (MacHardy, 1996; Vanneste, 2001, Way et al., 1991). To alleviate these production limitations, plant breeders strive to impart genetic resistance or resilience into improved cultivars. However, in apple, this process has been limited by breeding bottlenecks resulting in high interrelatedness of many cultivated varieties and breeding lines (Muranty et al., 2020; Migicovsky et al., 2021).

One approach to overcome this problem is to introduce novel genetics from crop wild relatives (CWRs). One successful example of CWR hybridization in apple was the introduction of resistance to apple scab caused by the fungus *Venturia inaequalis* (Gessler & Pertot, 2012). To that end, *M. floribunda* selection 821 was used to develop hybrids with *M.* × *domestica* which, ultimately, were used to identify a resistance locus named *Vf* (Hough et al., 1953; Williams et al., 1966). While these opportunities are afforded by the fact that *M.* × *domestica* readily hybridizes with multiple wild relatives, this process is limited by the considerable effort needed to purge undesirable traits and linkage drag from wild introgressions. This is in part due to the lack of genomic resources in wild species, the high heterozygosity due to self-incompatibility and the long generation time in apple (Migicovsky et al., 2021, Volk et al., 2015b, Sakurai et al., 2000).

Introgression breeding and genome editing, offer additional opportunities for further improvement of cultivated apples using genetics from wild species, and there has been a concerted effort, in recent years, to develop genomic resources in apple and its wild relatives (Velasco et al., 2010; Daccord et al., 2017; Chen et al., 2019; Zhang et al., 2019; Sun et al., 2020; Khan et al., 2022; Li et al., 2022). This has been bolstered by technological advancements that have enabled generating haplotype-resolved genomes of different progenitor species including *M. sieversii* and *M. sylvestris* (Sun et al., 2020). The *Malus* genus consists of 25 to 47 recognized species and additional hybrids (Robinson et al., 2001). Of those species, *M. fusca* and the other North American natives have been isolated from Asian and European gene pools used in the domestication of apple (Volk, 2019). Thus, *M. fusca* and its other North American relatives, offer untapped potential for unique disease resistance and abiotic tolerance traits.

As part of the effort to expand resources for breeding and genetics in apples, we report herein the high-quality, haplotype-resolved genome assembly of the *M. fusca* accession PI 589975. We used high-fidelity (HiFi) long reads, together with high-throughput chromosome conformation capture (Hi-C) to assemble and phase both haplotypes of this coastal Alaskan accession, which is resistant to fire blight, and moderately resistant to apple scab, as well as potentially other abiotic stresses (Way et al., 1991; Fiala 1994; USDA Agricultural Research Service 2015; Khan and Chao 2017; Papp et al., 2020; Dougherty et al., 2021). After gene annotation, we compared the synteny and genome architecture of this assembly to other domesticated and wild *Malus* genomes. Lastly, we explored synteny and presence-absence variance of candidate genes identified within the herein resolved fire blight resistance locus, *FB_MFu10* (Emeriewen et al., 2020), in comparison to other *Malus* genomes.

## Methods

### Plant Materials

For genomic DNA (gDNA) extractions, dormant scion cuttings of PI 589975 were obtained from the USDA Malus Collection at the United States Department of Agriculture (USDA) Plant Genetic Resources Unit (PGRU) located in Geneva, NY, USA. The cut ends of the dormant branches were placed into a beaker with a rehydration solution (Rose 100, Floralife, Waterboro, SC) using the manufacturer’s recommended concentration and placed under long-day conditions at 21°C. Once bud-break was achieved and expanded leaves reached 3 cm in length, the branches were transferred to dark conditions for 48 hours at 21°C. Leaves were then excised using a razor blade, weighed and split into 1 g samples. Tissue samples were immediately flash frozen in liquid N_2_ and stored at −80°C. Additional leaves was collected in early spring, when fresh growth was observed. This tissue was shipped overnight from the USDA Malus Collection, weighed and split into 0.25 g samples and immediately flash frozen in liquid N_2_ and stored at −80°C.

### DNA extraction

Genomic DNA (gDNA) and high-molecular weight (HMW) DNA were extracted from frozen dark-treated leaf tissues. For each extraction, replicates of 0.25 g of frozen leaf tissue were ground into a fine powder using a mortar and pestle chilled using liquid N_2_. For high-depth short-read whole-genome sequencing (WGS), gDNA was extracted using a Plant Pro DNA extraction kit (Qiagen, Germantown, MD) following the manufacturer’s protocol. Following extraction, DNA samples were cleaned and concentrated to improve quality using a Zymo Genomic DNA Clean & Concentration kit (Irvine, CA). Purified gDNA was checked for quality and quantity on a Nanodrop Spectrometer (Thermo Fisher Scientific, Waltham, MA) and Qubit 4 Fluorometer (Thermo Fisher Scientific, Waltham, MA). For the extraction of HMW DNA for long-read sequencing, Qiagen Genomic-tip 100/G kit was used following the protocol developed by Driguez et al. (2021). Extracted HMW DNA was checked for quality and quantity on a Nanodrop Spectrometer and Qubit 4 Fluorometer.

### Library Preparation and Sequencing

First, we obtained high-depth (∼100x) and high-quality short read sequence data. This sequencing of gDNA was carried out using an Illumina HiSeq platform (Illumina, San Diego, CA) with a read length of 150 bp in a paired end format. Library preparations and sequencing were performed by GeneWiz (South Plainfield, NJ). For PacBio (Menlo Park, CA) sequencing, we employed the HiFi protocol on a Sequel II system using a single SMRT cell. PacBio HiFi library preparation and sequencing was performed at the University of Maryland Institute of Genome Sciences (Baltimore, MD). Lastly, Hi-C libraries were prepared using the Phase Genomic Proximo Plant Kits (Seattle, WA) and sequenced on a MiSeq and HiSeq Illumina platform by GeneWiz.

### K-mer Based Genome Size Estimation

Adapters on the raw Illumina short-reads were removed by the sequencing provider, and duplicate reads were removed using nubeam-dedup software (Dai & Guan, 2020). To estimate genome size and heterozygosity, *k*-mers were counted using the Illumina WGS dataset with the software KMC 3 (v3.0) (Kokot et al., 2017) using the parameters ‘-k21 −t10 −m64 −ci1 - cs1000000’. We then exported the resulting *k*-mer count histogram into GenomeScope 2.0 following default instructions (Ranallo-Benavidez et al., 2020).

### Assembly and Scaffolding

HiFi FASTQ files were analyzed for adapter contamination using the HiFiAdapterFilt (v2.0.0) (Sims et al., 2022) and quality using FASTQC (v0.11.9) (Andrews, 2010). HiFi reads were then assembled using the HiFiasm (v0.16.1) assembler with the Hi-C paired-read option enabled to facilitate phasing of the contigs (Cheng et al., 2021). Once assembly was complete, each individual haplotype assembly was scaffolded using Chromonomer (v.1.13) with linkage map markers from *M. × domestica* (Bianco et al., 2014; Di Pierro et al., 2016; Catchen et al., 2020). Prior to scaffolding, marker sequences were aligned to the haplotype fastas using bwa mem (Li and Durbin 2009) and a pre-scaffold agp file was produced using the fasta2agp.py script from Chromonomer. The output agp files from Chromonomer were then converted to a Juicebox assembly file using Phase Genomic agp2assembly script (https://github.com/phasegenomics/juicebox_scripts). Assembly statistics were assessed using Merqury, Busco (v.5.3) using the embryophyta odb10 database, and gt seqstat (Rhie et al., 2020; Simão et al., 2015; Gremme et al., 2013).

Draft scaffolds were then validated by Hi-C contact matrixes. For this validation, Hi-C reads were mapped and filtered following the Arima Pipeline (https://github.com/ArimaGenomics/mapping_pipeline) (Ghurye et al., 2017). Mapped reads were then processed using Phase Genomic Juicebox utility Matlock (https://github.com/phasegenomics/matlock) and sorted following Phase Genomic suggested protocols. The sorted links file and Chromonomer assembly files were then passed through the 3D-DNA run-assembly-visualizer script to generate a Juicebox editable assembly file (https://github.com/aidenlab/3d-dna/blob/master/visualize/run-assembly-visualizer.sh). Any mis-joins or inversions in the scaffolds were then corrected in Juicebox (v1.11.08) by manual curation (Durand et al., 2016). Finalized scaffold assembly files were then used to generate a new representative FASTA using the contig sequences using the juicebox_assembly_converter script (https://github.com/phasegenomics/juicebox_scripts).

The haplotype FASTA were then aligned to GDDH13 v1.1 assembly using nucmer from the MUMmer package v4.0.0beta2 (Marçais et al., 2018), to assign scaffolds with known chromosome numbers. Chromosome05 was reverse-complemented to allow for easy comparisons between apple genomes. Plastid decontamination of the phased assemblies was done using the blobtools (v1.1.1) following the published protocols (Laetsch and Blaxter 2017). Additionally, during upload to the NCBI database, the system flagged ten scaffolds in haplotype 1 during the uploading of the genome as having additional mitochondrial contamination which were thus removed from the final assembly.

### Genome Annotation

The annotation of transposable elements (TE) for each genome was first conducted using EDTA (v1.9.6) (Ou et al., 2019) under default parameters except the ‘--species others’ option. In order to better compare repeat landscape contiguity across different *Malus* species (for the purpose of quality assessment) we annotated TEs, filtered, and consolidated the results using panEDTA (https://www.biorxiv.org/content/10.1101/2022.10.09.511471v1). The quality and contiguity of repeats was determined using the LTR Assembly Index (LAI) metric (Ou et al., 2018) from the LTR_retriever software (v2.9.0) (Ou and Jiang, 2018).

For annotation of the gene space, a comprehensive approach using MAKER2 was applied (Holt and Yandell 2011). Transcript evidence was assembled from publicly available RNA-seq libraries generated for a gene expression atlas (Rogers and van Nocker, 2013) of *Malus fusca* (NCBI Bioproject PRJNA267116). As this dataset contains more than 200 Gb of data, to provide EST evidence for annotation, 100 M pairs of reads were semi-randomly pulled from the BioProject’s corresponding SRAs using VARUS (Stanke et al., 2019). This approach ensures sufficiently high coverage across the genome while limiting data transfer. Reads were then mapped to each chromosome-only haplotype using STAR aligner with the options ‘--outSAMstrandField intronMotif’ and ‘--alignIntronMaxenabled 10 kb’ (Dobin et al., 2013). Transcripts for each haplotype were then assembled from the read alignments using StringTie2 (Kovaka et al., 2019).

FASTA sequences of the StringTie2 assembled transcripts were used as input into the first round of evidence-based annotation using MAKER2. Next, first round annotations were extracted using the extract_anno_evi.sh script and used to train *ab initio* gene predictors SNAP and Augustus (Korf 2004; Hoff and Stanke 2019). A second round of *ab initio* prediction was conducted using optimized transcripts from the first round of prediction and used as input into a final run of MAKER2. Annotations of rRNA and tRNA features used RNAmmer and tRNAscan-SE with default options (Lagesen et al., 2007; Chan and Lowe 2019).

### Synteny analysis

Comparison of gene synteny between the haplotype 1 assembly and two wild species (*M. sieversii* and *M. sylvestris,* Sun et al., 2020) as well as three *M. × domestica* genomes (GDDH13 v1.1, Gala, and Honeycrisp; Daccord et al., 2017; Sun et al., 2020; Khan et al., 2022) was performed with the Python MCScanX pipeline v1.1.12 (Tang et al., 2008; Wang et al., 2012). Briefly, annotation gff files were downloaded from the Genome Database for Rosaceae (GDR) (Jung et al., 2019) and converted to bed format using jcvi.formats.gff. A pairwise synteny search was performed. To mitigate the impact of the recent whole genome duplication in *Malus*, only the reciprocal best hits (‘--cscore=0.99’) were used for establishing the high-quality synteny blocks utilized in syntenic depth comparisons and plotting of karyotypes and macrosynteny and microsynteny plots, as well as syntenic block depths.

### Haplotypic structural variation

SVs between the two haplotypes were detected by aligning the two FASTA files using nucmer with the settings ‘--maxmatch −l 100 −c 500’ (Marçais et al., 2018). The resulting gziped delta file was uploaded to the Assemblytics web interface (http://www.assemblytics.com/) for analysis using a maximum variant size of 50 kbp (Nattestad and Schatz 2016). The results were exported as a bed file and imported into R for plotting. Deletions overlapping genes and exons were identified using Bedtools intersect (Quinlan and Hall 2010). Haplotypic deletions were validated by mapping the short Illumina reads. First, reads were aligned to the haplotype 1 assembly using bwa mem with default settings (Li and Durbin 2009). The average coverage for deletions was then compared to random non-deletion regions. Mean coverage across each deletion was calculated by Bedtools ‘coverage −mean’ for each predicted deletion. Bedtools shuffle was used to collect 100x random non-deletion regions of the same size of each deletion. The coverage distributions of deletion and non-deletion regions were compared using a 1000x bootstrapped Kolmogorov–Smirnov test from the ‘matching’ R package (Sekhon, 2011). The sorted bam file was then loaded into samplot (Belyeu et al., 2021) to visualize selected deletions, and plot the read coverage and identify discordant mapping and long insert sizes.

### Synteny at the FB_Mfu10 resistance locus

To identify the location of the previously reported fire blight resistance locus, the primers for the markers flanking the fine-mapped region of fire blight resistance on chromosome 10 were extracted from Emeriewen et al., (2018) and their sequence was aligned to haplotype 1 using BLASTN. Specifically, markers FR39G5T7xT7y and FR46H22 which were shown to delimit the locus to 0.33 cM were used to locate the narrowest region. As multiple BLAST hits were reported, the location was also confirmed by BLASTN of a candidate gene reported by Emerienwen et al., (2021), however the broader region between the markers was considered for further analysis. The region was then interrogated for microsynteny between the *M. fusca* haplotype 1 and the respective regions on chromosome 10 from the susceptible *M. × domestica* genomes, including GDDH13 v1.1 (Daccord et al., 2010), Gala (Sun et al., 2020), Honeycrisp (Khan et al., 2022), *M. sieversii* and *M. sylvestris* (Sun et al., 2020). Because MCScanX (Wang et al., 2012) excludes most tandem gene arrays in synteny analysis due to the difficulty assigning true matches in these arrays, matches between candidate genes in the region were analyzed in a phylogenetic manner. Peptide sequences were extracted from each ortholog candidate gene and aligned using Custal Omega within the ETE3 toolkit (v.3.1.2) (Huerta-Cepas et al., 2016; Sievers and Higgins 2018). ETE3 then executed the standard FastTree workflow for generation of a phylogenetic tree (Price et al., 2010). Candidate gene orthologs *Msy10g019590*, *Mdg10g019850*, and *MD10G1207300* from *M. sylvestris,* Gala, and GDDH13, respectively, were removed from the analysis due to potential mis-joined annotations or fragmented structure/pseudogene identity to facilitate more clear phylogenetic clade membership. Syntenic orthologs were then classified by the clade membership.

### Data Availability

The raw sequence data generated from this project can be retrieved from NCBI SRA database under BioSample SAMN31658743 and phased assemblies under BioProjects PRJNA899490 and PRJNA899491. Genome assemblies and annotation files can be retrieved from the GDR accession number tfGDR1066. Code for producing figures for the manuscript is available at github.com/bmansfeld/mfusca_figs/.

## Results and discussion

The *M. fusca* accession we chose to sequence was PI 589975 (GMAL 2891) from the USDA Germplasm collection (Fig 1A-C) (USDA Agricultural Research Service, 2015). PI 589975 was originally sampled from Scow Bay near Petersburg, Alaska near the edge of woods along the beach (Fig 1D). The accession was donated to the USDA collection in August of 1988 by Michael Medalen and is a member of the core collection grown on ‘Budagovsky 9’ rootstock (USDA Agricultural Research Service, 2015).

**Figure 1.**
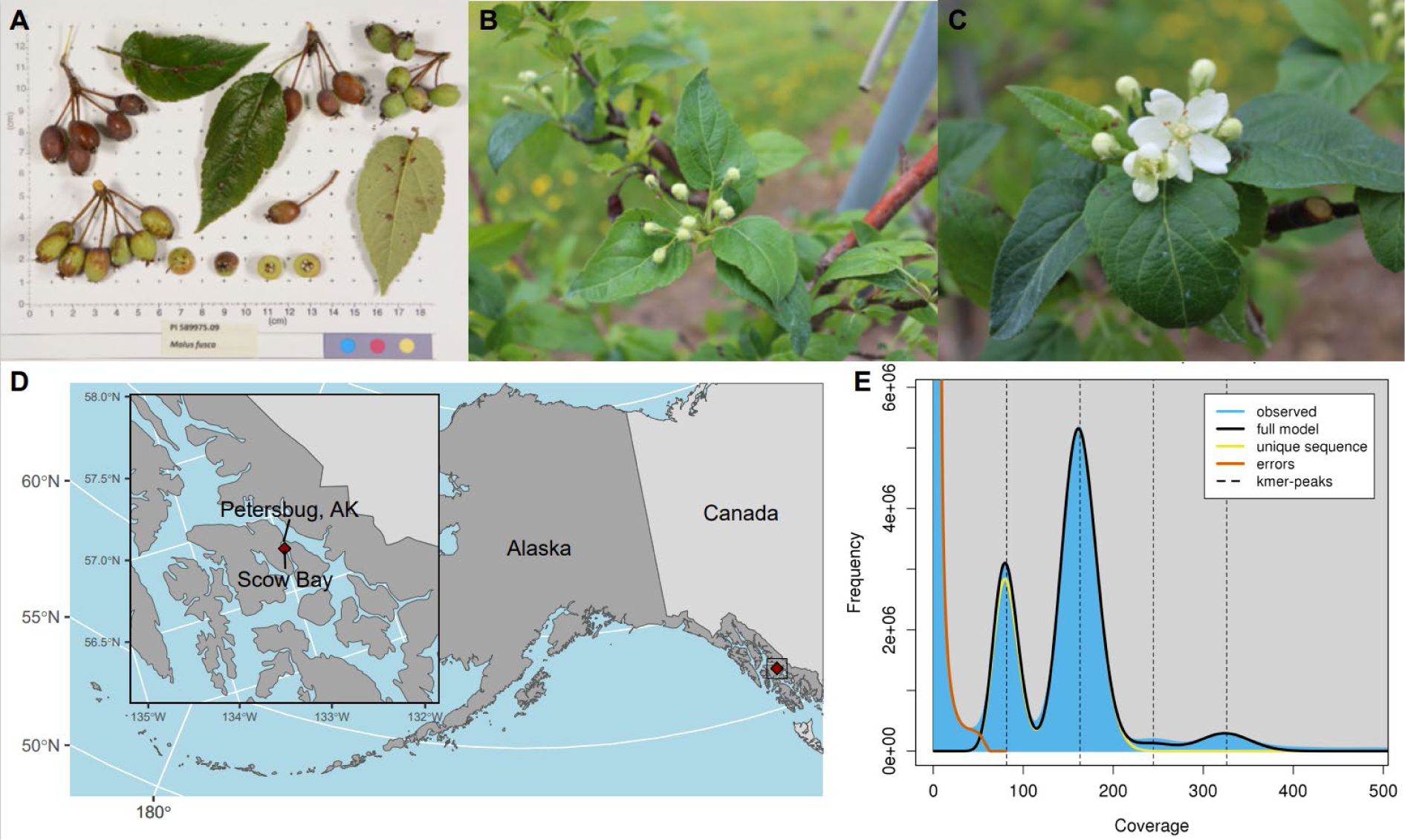
**A)** Malus fusca accession PI 589975 GRIN-Global identification photograph. **B)** Ballon stage blossoms of the living PI 589975 accession in the USDA Plant Germplasm Repository Unit, Geneva, NY. **C)** King bloom stage of the living PI 589975 accession in the USDA Plant Germplasm Repository Unit, Geneva, NY. **D)** Map of the approximate collection position of the PI 589975 in Alaska. Insert is of the specific location of Scow Bay near Petersburg, AK. **E)** *k*-mer count plot from GenomeScope. The haploid genome size was predicted to be 694 Mb with approximately 56% repetitive sequence and an estimated heterozygosity of 0.8%.

### Reference quality genome assembly

We obtained 95 Gb and 21.7 Gb of sequence for this accession of *M. fusca* from Illumina and PacBio HiFi sequencing, respectively. The Illumina sequencing based *k*-mer analysis yielded an estimated genome size of 694 Mb and heterozygosity of 0.8% (Fig 1E). Thus, the HiFi sequencing represented 31.3x coverage from 1.4 million reads, which is sufficient for *de novo* haplotype resolved assembly using HiFiasm (Cheng et al., 2021, Cheng et al., 2022). High quality Hi-C libraries for *M. fusca* were extremely difficult to generate and after several attempts with different kits were still of consistently of low concentrations, suggesting issues with extraction and amplification of the proximal DNA contacts. Regardless, we sequenced one Hi-C library and obtained ∼130x data with sufficient quality for haplotype phasing. HiFiasm successfully resolved the two haplotype assemblies of 682 and 644 Mb in length (SFig 1) and contig N50 of 18.7 and 21.4 Mb, respectively (Table 1). Some contigs were near chromosome length, with the largest contig assembled at 42.3 Mb.

**Table 1.** Genome assembly statistics.

| Assembly | Length (bp) | Number of contigs | N50 | Contigs >1Mb | % of estimated size | Merqury QV | Merqury QV error rate | Merqury *k*-mer completeness | % BUSCO | Complete single copy Busco | Complete duplicate Busco | Fragmented Busco | Missing Busco |
| --- | --- | --- | --- | --- | --- | --- | --- | --- | --- | --- | --- | --- | --- |
| *HiFi Contigs* |  |  |  |  |  |  |  |  |  |  |  |  |  |
| Haplotype 1 | 682,253,989 | 1,141 | 18,869,149 | 48 | 98.31% | 62.39 | 5.77E-07 | 86.82% |  |  |  |  |  |
| Haplotype 2 | 644,255,151 | 273 | 21,444,431 | 41 | 92.83% | 67.14 | 1.93E-07 | 86.51% |  |  |  |  |  |
| Combined | 1,326,509,140 | 1,414 |  |  |  | 64.09 | 3.90E-07 | 98.21% |  |  |  |  |  |
| *Scaffolds* |  |  |  |  |  |  |  |  |  |  |  |  |  |
| Haplotype 1 | 682,257,189 | 1,109 | 36,508,756 | 18 | 98.31% |  |  |  |  |  |  |  |  |
| Haplotype 2 | 644,257,951 | 245 | 36,111,528 | 17 | 92.84% |  |  |  |  |  |  |  |  |
| *Decontaminated Assembly* |  |  |  |  |  |  |  |  |  |  |  |  |  |
| Haplotype 1 | 651,182,355 | 235 | 36,779,757 | 18 | 93.83% |  |  |  | 98.70% | 993 | 600 | 11 | 10 |
| Haplotype 2 | 637,756,398 | 97 | 36,111,528 | 17 | 91.90% |  |  |  | 98.82% | 992 | 603 | 11 | 8 |
| *Chromosome-only Assembly* |  |  |  |  |  |  |  |  |  |  |  |  |  |
| Haplotype 1 | 637,459,684 | 17 | 36,779,757 | 17 | 91.86% |  |  |  | 98.76% | 996 | 598 | 11 | 9 |
| Haplotype 2 | 631,922,950 | 17 | 36,111,528 | 17 | 91.06% |  |  |  | 98.88% | 994 | 602 | 11 | 7 |

As the assemblies were extremely contiguous, we leveraged the high relatedness of *M. fusca* to *M. × domestica* (Robinson et al., 2001; Nikiforova et al., 2013; Volk et al., 2015) and scaffolded both haplotypes using a *Malus* combined linkage map (Bianco et al., 2014; Di Pierro et al., 2016; Catchen et al., 2020). We placed the contigs into 17 chromosome sized pseudomolecules (Fig 2A). As an orthogonal verification of scaffolding, we inspected contact maps using Hi-C alignments with Juicebox (Durand et al., 2016). The Hi-C contact maps showed high proximal interactions with the scaffolds in contig placement and order, and needed minor manual curation (SFig 2A,B).

**Figure 2.**
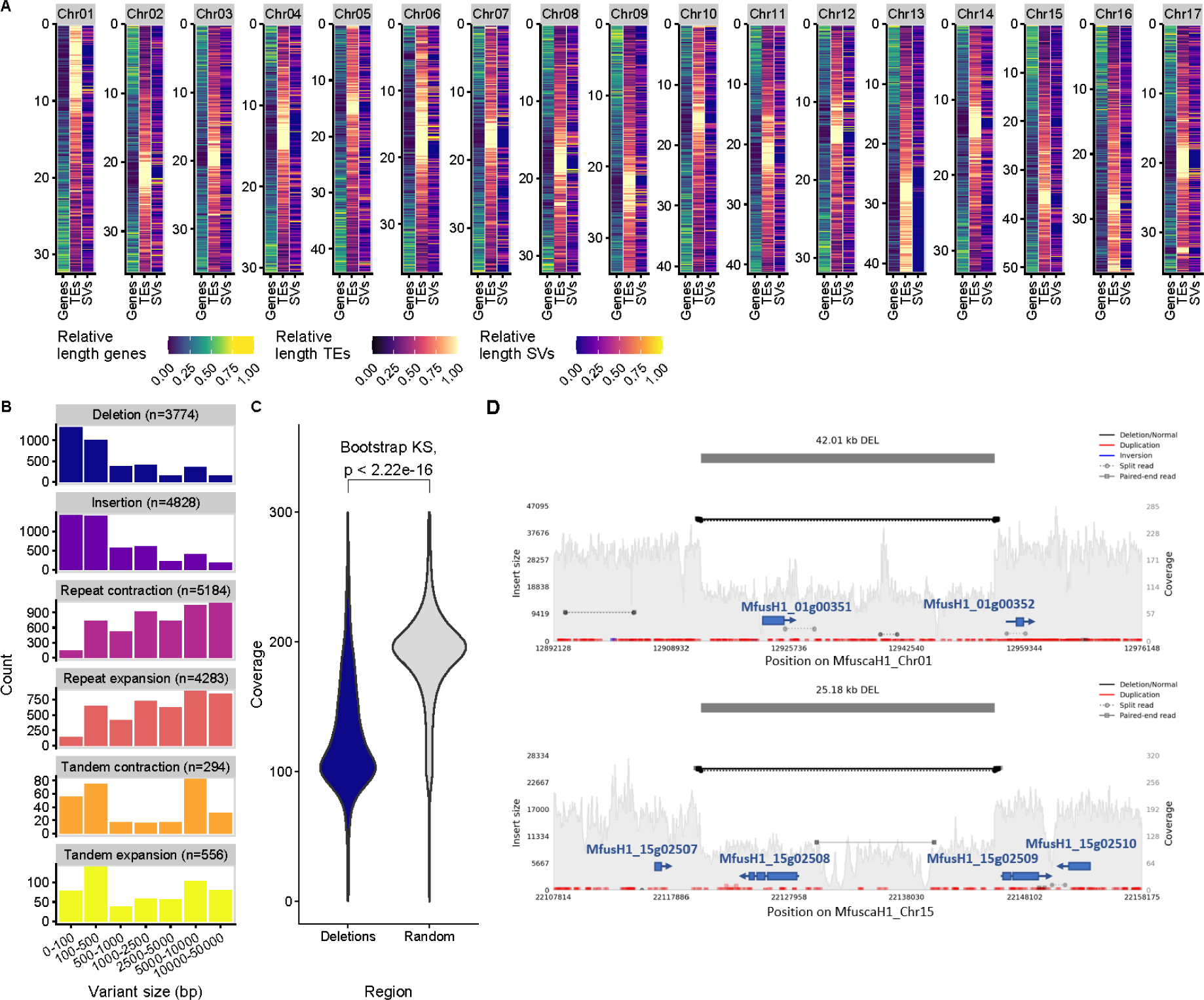
*Malus fusca* genome assembly annotation and haplotype comparisons. **A)** Distribution heat maps of annotated features across all 17 chromosomes (genes, TEs-Transposable Elements, and SVs-Structural Variations). The relative cumulative length of each feature type was calculated within a 100 Kbp window. **B)** Size distribution of different haplotypic SV within the two haplotypes. **C)** WGS read coverage between deletions vs random positions within the genome. The difference in distance distributions was evaluated by a 1000x bootstrapped Kolmogorov–Smirnov (KS) test. **D)** Examples of large haplotypic deletions that contain annotated genes within the SV, which result in hemizygous genes. Grey histograms represent depth of coverage (right y-axis) of short reads mapped to the haplotype 1 assembly. Reads pairs spanning the identified deletions are highlighted in black and the size of the insert between pairs is denoted by the position vs. the left y-axis. Reads paired with dotted lines indicate split reads that map to both sides of the deletion. Read pairs in red indicate sequence duplications. Gene models are denoted in blue.

The two haplotype assemblies (or haplomes) are 94% and 91% complete based on the estimated genome size. Merqury *k*-mer based analysis showed a completeness score of 98.2% for the diploid assembly and a QV score of 64.1, the highest to date in any wild *Malus* species. We report a complete genome BUSCO (Simão et al., 2015) score of 98.8 and 98.9% for the two assemblies respectively. The duplicate BUSCO score was 37.2 and 37.4%, comparable to other *Malus* spp. (Sun et al., 2020), and indicative of the relatively recent paleo-duplication event in *Malus* (Velasco et al., 2010). In summary, after scaffolding and decontamination, we obtained two haplotype assembles, which were highly contiguous, nearly complete, and extremely accurate with lengths of 651 Mb and 634 Mb and with scaffold N50s of 36.8 and 36.1 Mb, respectively (Table 1).

To further evaluate the assembly contiguity of repetitive sequences, we computed the LAI metric (Ou et al., 2018) of *M. fusca* haplotypes and other published apple genomes/haplotypes (Velasco et al., 2010; Daccord et al., 2017; Chen et al., 2019; Linsmith et al., 2019; Zhang et al., 2019; Sun et al., 2020; Khan et al., 2022; Li et al., 2022) (STable 1). Our phased haplotypes have the highest LAI scores and are greater than 20 (“gold” standard based on Ou et al., 2018), indicating that repeats and TEs were assembled with high quality (STable 1). The use of HiFi reads allowed us to sequence through repeats accurately and enabled assembly of chromosome-scale contigs. As a result, our assemblies represent one of the highest quality diploid pome fruit genome published to date.

### Genome Annotation

The availability of multiple *Malus* genomes allowed side-by-side comparisons of their repeat content and contiguity but required consistent TE annotation for this purpose. We collected all published *Malus* genomes and the European pear (*Pyrus Communis*) genome (Velasco et al., 2010; Daccord et al., 2017; Chen et al., 2019; Linsmith et al., 2019; Zhang et al., 2019; Sun et al., 2020; Khan et al., 2022; Li et al., 2022). Together with the two haplotypes of *M. fusca* genome, we created a TE annotation for the *Malus* genus with the European pear genome as an outgroup. We found that the *Malus* genome assemblies contained between 48.6% to 62.43% TEs, while the European pear genome contained roughly 45.3% (STable 1). The two *M. fusca* assemblies fell within this range with an average TE content of 55% (Table 2). Long Terminal Repeat (LTRs) retrotransposons were the most abundant TE within these genomes. The two *M. fusca* haplotypes were found to contain 351.68 and 346.65 Mb of annotated transposable elements (TE), equating to 55.17% and 54.86%, respectively, of each haplotype’s total length as predicted by the *k-*mer-based approach (Fig 1E, Table 2).

**Table 2.** Genome annotation statistics.

| Assembly | Haplotype 1 | Haplotype 2 | Combined |
| --- | --- | --- | --- |
| *Intergenic* |  |  |  |
| LTR - Copia | 11.69% | 11.67% |  |
| LTR - Gypsy | 15.41% | 15.11% |  |
| LTR - Unknown | 16.75% | 16.72% |  |
| TIR-CACTA | 1.25% | 1.25% |  |
| TIR-Mutator | 4.54% | 4.53% |  |
| TIR-PIF Harbinger | 2.39% | 2.40% |  |
| TIR-Tc1 Mariner | 0.03% | 0.03% |  |
| TIR-hAT | 2.80% | 2.81% |  |
| Low complexity | 0.02% | 0.02% |  |
| nonTIR - helitron | 0.29% | 0.31% |  |
| Total % | 55.17% | 54.86% |  |
| Total bp | 351.68 Mb | 346.65 Mb |  |
| *Genic* |  |  |  |
| % Complete BUSCO genes | 97.5 | 96.6 |  |
| Complete single copy BUSCO genes | 1006 | 1010 |  |
| Complete duplicate BUSCO genes | 557 | 549 |  |
| Fragmented BUSCO genes | 14 | 13 |  |
| Missing BUSCO genes | 37 | 42 |  |
| Genes | 46,622 | 45,853 | 92,475 |
| tRNA genes | 475 | 472 | 947 |
| rRNA gene | 673 | 822 | 1495 |

Footnote: BUSCO (v.5.3) using the embrophyta odb 10 *n*=1614

Previous exhaustive sequencing of the *M. fusca* transcriptome (PRJNA267116), including 72 different tissue types and developmental stages, allowed us an unprecedented opportunity to thoroughly annotate the genome of *M. fusca*. In total, we annotated 46,622 and 45,853 genes for each of the haplotype assemblies, respectively (Table 2). Roughly 75% of annotated genes had an Annotation Edit Distance of less than 0.25 indicating high support by transcriptional and protein evidence (SFig 3). A BUSCO analysis of the two annotated transcriptomes indicated 96.8% and 96.6% completeness, respectively. Additionally, we annotated 947 tRNA and 1495 rRNA genes between the two haplotypes (Table 2). These resources amount to one of the most thoroughly annotated apple genomes to date (Fig. 2A).

**Figure 3.**
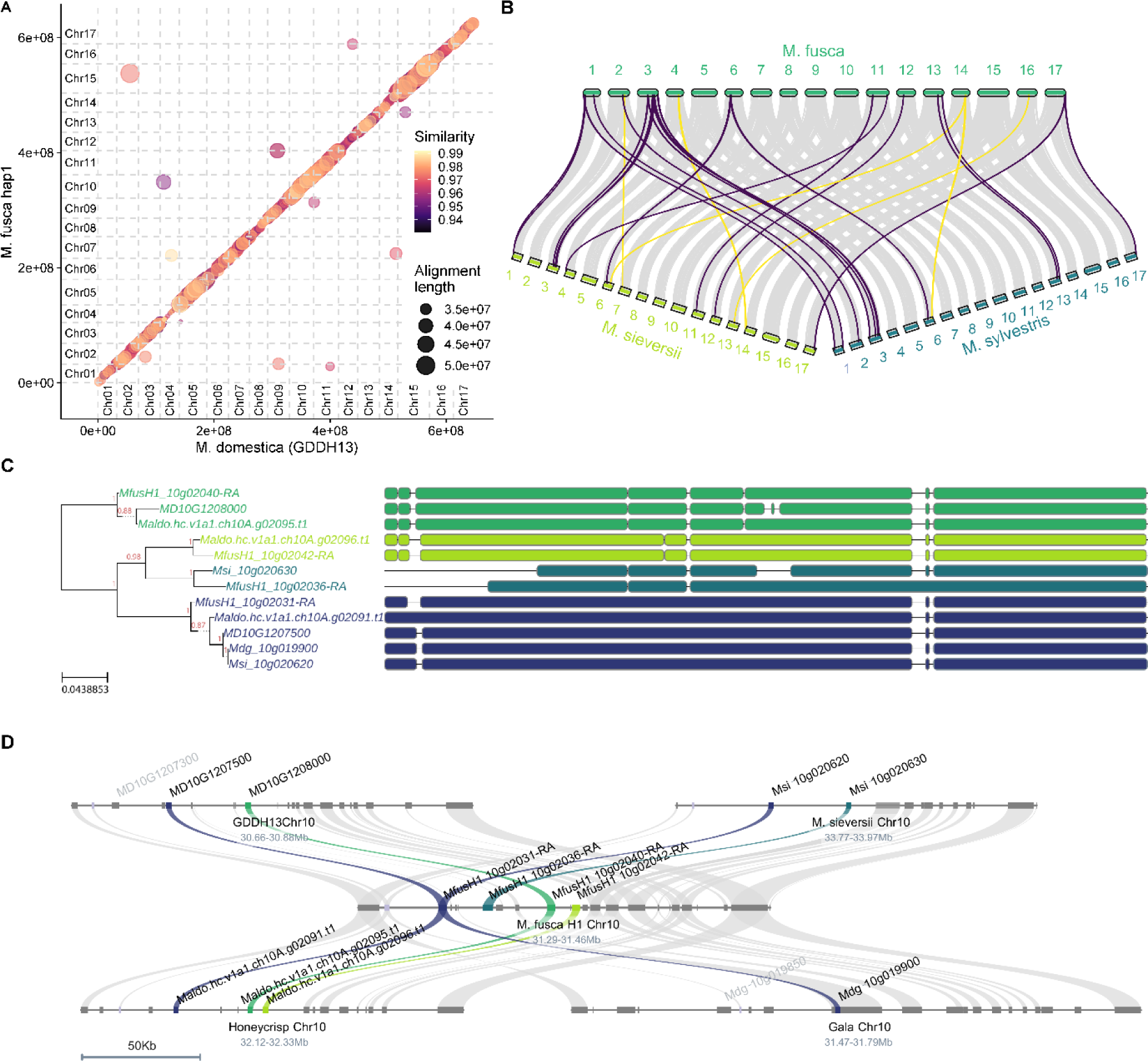
Comparative genomics within *Malus* and the *FB_Mfu10* fire blight resistance locus. **A)** Alignment length and sequence similarity between *M. fusca* haplotype 1 and *M. × domestica* GDDH13. **B)** Gene synteny between the *M. fusca* genome and wild apple species *M. sieversii* and *M. sylvestris* genomes (purple = inversions, yellow = translocations). **C)** Phylogenetic tree and peptide alignments of genes encoding G-type lectin S-receptor-like serine/threonine-protein kinases within the *FB_Mfu10* locus in *M. fusca, M. × domestica* cultivars, and *M. sieversii.* **D)** *Microsynteny at the Mfu10 locus*. Copy number variation of G-type lectin S-receptor-like genes within the *FB_Mfu10* locus associated with fire blight resistance in *M. fusca* (center) compared to susceptible *M. sieversii* (top right), and *M.* x *domestica* Golden Delicious (top left), *Honeycrisp* (bottom left), and Gala (bottom right). Orthologs of G-type lectin S-receptor-like serine/threonine-protein kinase genes are annotated by locus id and syntenic genes are annotated by color between the four genomes. Gene colors in panels C and D match to show the phylogeny based synteny inference. Gene fragments with sequence similarity to these receptors are also denoted but shaded in light grey.

### Haplotypic polymorphisms and structural variation in M. fusca

We aligned the two haplotype assemblies against each other and identified SNPs and large SVs between the two haplotypes. We found a total of 2,454,873 SNPs and 1,969,639 small InDels. However, high heterozygosity in plants manifests not only as single nucleotide polymorphisms, and recent work in other heterozygous crop species (*e.g.*, Zhou et al., 2019, Mansfeld et al., 2021) revealed that large haplotypic structural variation (SV) contributes to differences between the haplotypes in these species. Our highly contiguous *M. fusca* haplotype assemblies thus present an opportunity to evaluate the variation between haplotypes in an outcrossing wild relative of a cultivated fruit tree. Apart from the small haplotypic differences, we identified over 18,000 large SVs ranging in size from 50-50,000 bp (SFile 1). These included large insertions, deletions, tandem duplications/contractions, and repeat expansions/contractions (Fig. 2B). These SVs impacted greater than 77 Mb of total genomic space. For comparison to another highly heterozygous crop genome analyzed by similar methods, the volume of haplotypic SVs detected in this assembly was larger than that observed in the African cassava genome (Mansfeld et al., 2021), even though heterozygosity in cassava (∼1.4%) was estimated at nearly double that of *M. fusca*. This is likely due to the high completeness, contiguity, and scaffolded nature of both haplotypes in the *M. fusca* assembly, which allows for more accurate and thorough haplotypic comparisons. This suggests that modern haplotype resolved assembly strategies (Cheng et al., 2021) such as the one used herein, have crucial implications for the ability to detect these important haplotypic SVs.

We were especially interested in the 3,774 large haplotypic deletions observed, as these might cause gene hemizygosity (*e.g.*, Zhou et al., 2019) or impact important gene function (STable 2). To further validate these large InDels, we compared short read coverage at the deletion sites to random (non-deletion) regions of the same size (Fig. 2C). As expected, we found that mapped read coverage in the deletions was roughly 50% of, and significantly different from that of randomly selected genome regions (1000x bootstrap KS test, p-value < 2.22e-16), suggesting that most haplotypic deletions identified by sequence alignment are accurate. We further explored specific examples of large deletions that overlapped with genes and further validated these cases by examining inflated insert sizes between paired reads. Overall, we identified 3,853 unique cases where exons overlapped with a haplotypic deletion (Fig. 2D; STable 3). This included instances of complete hemizygosity of some genes due to these large deletions. For example, a 42kb heterozygous deletion on chromosome 01 causes hemizygosity of *MfusH1_01g00351*, while a second 25kb deletion on chromosome 15, removes one allele of *MfusH1_15g02509*. Similar structural variation has been shown to have substantive effects on important agricultural traits. For example, berry color in Chardonnay grape is likely altered due to hemizygosity at the *MybA* locus (Zhou et al., 2019). The above two validated deletions also remove the entire up-stream regions for *MfusH1_01g00352* and *MfusH1_15g02509*, respectively. Large haplotypic deletions have been also implicated in impacting gene expression profiles by modifying their cis-regulatory landscape (Sun et al., 2020; Mansfeld et al., 2021), as well as have important consequences for the epigenomic landscape (Zhong et al., 2022). Using this assembly, gene hemizygosity can now be taken into account when attempting to introgress traits from *M. fusca* in breeding efforts. Future research should thus explore the impacts of these large haplotypic SVs on allele specific expression in regard to important traits in *M. fusca* and how these might be useful to breeders.

### Comparing apples to apples: synteny within Malus

The emergence of 3^rd^ generation sequencing technologies and improved scaffolding methods *(e.g*., Hi-C and Omni-C sequencing), have resulted in numerous high-quality apple genomes to compare against. The first, high-quality apple genome developed using long-reads was GDDH13, a doubled haploid of Golden Delicious, and it serves as an inbred reference genome (Daccord et al., 2017). We aligned *M. fusca* haplotype 1 to GDDH13 to identify regions of high sequencing similarity and length. Even though *M. fusca* has been kept separate from the rest of the domestication history of apple, we still observe high sequence similarity and co-linearity between *M. fusca* and *M. × domestica* (Fig. 3A). We identified 2,474,476 SNPs and 1,400,493 small indels as well as 17,467 large SVs (affecting 114 Mbp) between the species (SFig 4, Sfile 2). Similar species-specific variations have been shown to underlie useful crop improvement traits. In tomato for example, SV between wild and domesticated material was shown to impact fruit size and volatile composition (Alonge et al., 2020). Thus, the genome assembly herein will support future work to establish the role of SV on similar traits in *Malus*.

We also performed gene synteny analysis with two wild progenitors of domesticated apple, *M. sieversii* and *M. sylvestris* (Sun et al., 2020). Comparisons of chloroplast genomes indicate that *M. fusca* is closely related to these species, which suggests Asiatic origins for *M. fusca* (Robinson et al., 2001; Nikiforova et al., 2013; Volk et al., 2015). Thus perhaps, these wild species were only recently separated geographically by the submersion of the Beringia land bridge, that connected Asia to North America (Williams 1982; Routson et al., 2012). Indeed, we observed high syntenic relationships between the three species (Fig. 3B). However, several macro-scale variations were found, including large inversions and translocations. Most of the variation from *M. fusca* was shared by the other wild species, however some species-specific translocations and inversions were observed. For example, several inversions were observed on chromosome 03 that were shared between *M. fusca* and the two other wild apples, but the translocation between *M. fusca* chromosome 16 and *M. sieversii* chromosome 13 was specific to that comparison. Taken together, the relatively high whole-genome synteny and limited macro-variations support the Asiatic origins of *M. fusca* (Robinson et al., 2001; Nikiforova et al., 2013; Volk et al., 2015). More population genetic work, focused on whole genome evolution, should be performed in the future to better understand the relationship between these wild species. It will be interesting to explore how selection and cultivation of *M. fusca* by indigenous people in the Pacific Northwest of North America (Wyllie de Echeverria 2013), and its isolation from the domestication history of *M. × domestica*, have impacted traits that may be utilized in future improvement of apple cultivars and rootstocks.

### Resolving the Fire blight Resistance Locus (FB_Mfu10)

Apart from the unique climate and temperature cline that *M. fusca* has adapted to (Routson et al., 2012), the geographic localization of *M. fusca* in North America has potentially allowed for important co-evolution with *Erwinia amylovora.* This native North American bacterium causes the disease fire blight and is the most important constraint on pome production in the world (Norelli et al., 2003; Van der Zwet et al., 2012). Importantly, most *M. fusca* accessions are tolerant or resistant to fire blight (Dougherty et al., 2021), and indeed a noted resistance locus was identified on *M. fusca* chromosome 10 through screening of segregating populations derived from crosses of *M. fusca* with the susceptible *M.* x *domestica* cultivar Idared (Emeriewen et al., 2017). In that research, a genetic map was developed positioning a QTL on Chromosome 10 (*FB_Mfu10*) that explained 66% of the variation (Emeriewen et al., 2017; Emeriewen et al., 2020). Further analysis of that region using Illumina-and Nanopore-sequenced BACs, identified a potential candidate gene, as well as other repetitive fragments of high sequence similarity to the candidate gene (Emeriewen et al., 2018; Emeriewen et al., 2022). We sought to leverage our highly contiguous assembly of this fire blight resistant *M. fusca* accession to help in resolving this important locus and further analyze the genes therein, which likely contribute to this resistance trait.

Within the fine-mapped boundaries identified by Emeriewen et al., (2018) we identified several genes including a tandem duplication array consisting of four copies (*MfusH1_10g02031, MfusH1_10g02036, MfusH1_10g02040,* and *MfusH1_10g02042*) of the G-type lectin S-receptor-like serine/threonine-protein kinase genes implicated by Emeriewen et al., (2018;2022). Since similar resistance receptors (*i.e.,* R-genes) are often part of such tandem duplications; we hypothesized that apart from sequence polymorphism, copy number variation (CNV) within this locus could potentially contribute to the resistance phenotype. We thus performed micro-synteny level comparisons between the resistant *M. fusca* and three susceptible *M.* x *domestica* cultivars (Honeycrisp, Golden Delicious, and Gala) (Daccord et al., 2017; Sun et al., 2020; Khan et al., 2022). We observed CNV in genes encoding these receptor-like genes that correlated with reported resistance phenotypes. While the resistant *M. fusca* carries four copies of this R-genes, ‘Honeycrisp’ contains three, ‘Golden Delicious’ has two full and one fragmented gene, and finally, ’Galà contains two copies of which one is a truncated fragment (Fig. 3D). ’Galà is moderately to highly susceptible, ’Honeycrisp’ is moderately susceptible to moderately resistant, and ’Golden Delicious’ is moderately resistant (Kostick et al., 2019; Dougherty et al., 2021). However, it should be noted that the GDDH13 assembly is of a doubled haploid of ’Golden Delicious’ and thus only represents the haplotype of this cultivar.

We expanded the comparative genomics analysis of *FB_Mfu10* locus to compare *M. fusca* to the other sequenced wild *Malus* relatives – *M. sieversii* and *M. sylvestris* (Sun et al., 2020). The accessions of *M. sieversii* (PI 613981) and *M. sylvestris* (PI 633825) are reportedly susceptible and resistant to fire blight, respectively, presenting an ideal opportunity to test our hypothesis (USDA Agricultural Research Service, 2015; B. Gutierrez personal communication). *M. sieversii* was found to have two copies of the G-type receptor gene within the locus while *M. sylvestris* had one large (>8000 bp) ortholog annotated, likely representing three copies misjoined as one gene (SFig 5). This result lends support to our hypothesis that CNV correlates with the resistance phenotype. However, *M. sylvestris* may only contain three copies which are similar to the susceptible Honeycrisp and Golden Delicious alleles. Thus, sequence variation that affects gene function or expression may also contribute to the resistance phenotype. This type of variation is evident in the sequence alignment and phylogenetic relationship between the genes (Fig 3C). Alternatively, *M. sylvestris* might have other loci contributing to resistance elsewhere in the genome. Similar examples of CNV of R-genes have been reported in maize (Chavan et al., 2015), soybean (Cook et al., 2012; Lee et al., 2015), and R-genes were found to be enriched for CNV in the genome of *M.* x *domestica* (Boocock et al., 2015). Additionally, Linkage Group 10 has been previously implicated in resistance to fire blight within a mapping population of *M.* x *domestica* generated from ‘Florina’ x ‘Nova Easygro’ (Le Roux et al., 2010), suggesting some contribution of genes on this chromosome already within the *M.* x *domestica* germplasm.

Previously, Fahrentrapp et al., (2013) and Emeriewen et al., (2018) speculated that the single candidate genes that underlie fire blight resistance loci in *Malus spp.* are not products of co-evolution with the *Erwinia* due to their lack of positioning within clusters of paralogs (*i.e.*, arrays). However, the results presented by Emeriewen et al., (2022) and our genome support a co-evolutionary origin of the fire blight resistance in *M. fusca*. Our assembly demonstrates that the Emeriewen et al., (2022) candidate R-gene was located within a tandem array of similar R-genes, which also span into other *Malus spp.* and domesticated cultivars. Taken together, it can be hypothesized that the *FB_Mfu10* locus underwent selection for increased CNV of the R-gene in response to co-evolution with *Erwinia* in their overlapping habitats. Furthermore, it can be hypothesized that other North American species of *Malus* (*M. angustifolia, M. coronaria*, and *M. ioensis*) co-evolved even stronger resistance due to a longer evolutionary history with *Erwinia* in the pathogen’s center of origin in the eastern North America (van der Zwet 2006; Stukenbrock and McDonald 2008; McGhee and Sundin 2012; Zeng et al., 2017). In support of this hypothesis, *M. angustifolia* was found to exhibit lower susceptibility of natural infection by *Erwinia* than *M. fusca* (Dougherty et al., 2021). Exploring this locus in other native North American *Malus* species should help shed light on this possibility and help identify crucial alleles that confer higher and more durable resistance to infection.

## Conclusion

The herein described haplotype-resolved genome and annotation of *M. fusca* will add to the many recent developments in *Malus* genomics. Moreover, it provides an example of the opportunity that is afforded by increased access to long-read sequencing and improved genome assembly, scaffolding, and annotation methods in generating high-quality genomes for CWRs. Importantly for this work, our high-quality genome provides a valuable resource for understanding the genetic basis of important traits in this species and in the genus at large, such as disease resistance and stress tolerance, which are crucial for apple breeding programs moving forward. Furthermore, this study also highlights the significance of preserving wild apple relatives as a source of genetic diversity for future breeding efforts.

## Supporting information

STable 1

Stable 2

STable 3

SFile 1

SFile 2

## Acknowledgements

The authors acknowledge Dr. Becky Bart at the Donald Danforth Plant Science Center for donation of computational time and Dr. Chris Dardick for critical input on the manuscript. Additionally, the authors thank Dr. Sean Rogers for his efforts in developing the transcriptomic resources used for annotation.

## Supplemental Materials

**Supplemental Figure 1.** Merqury *k*-mer multiplicity analysis of the phased chromosome-only assembly.

**Supplemental Figure 2.** Hi-C contact maps with the scaffolded genomes. A) Haplotype 1. B) Haplotype 2.

**Supplemental Figure 3.** Annotation Edit Distance (AED) of the gene annotation for Haplome 1 and 2.

**Supplemental Figure 4.** Assemblytics structural variation between *M. fusca* and *M.* x *domestica* GDDH13.

**Supplemental Figure 5.** Synteny within the *Mfu10* locus associated with fire blight resistance in *M. fusca* (top) compared to *M. sylvestris* (bottom). Orthologs of G-type lectin S-receptor-like serine/threonine-protein kinases genes (R-genes) are annotated by locus id and syntenic genes are annotated by color between the four genomes.

STable 1. TE annotations and LTR assembly index (LAI) scores of *Malus* and *Pyrus* genomes.

STable 2. Structural variant deletions between *M. fusca* haplotype 1 and 2.

STable 3. Structural variants deletions between *M. fusca* haplotype 1 and 2 that overlap with exons.

SFile 1. *M. fusca* Haplotype 1 vs *M. fusca* Haplotype 2 structural variation BED file.

SFile 2. *M. fusca* Haplotype 1 vs GDDH13v1.1 structural variation BED file.

## Notes

### Competing Interest Statement

The authors have declared no competing interest.

