## Supplemental Figures for "A haplotype resolved chromosome-scale assembly of North American wild apple *Malus fusca* and comparative genomics of the fire blight *Mfu10* locus. Genome of the Pacific Crabapple *Malus fusca*"

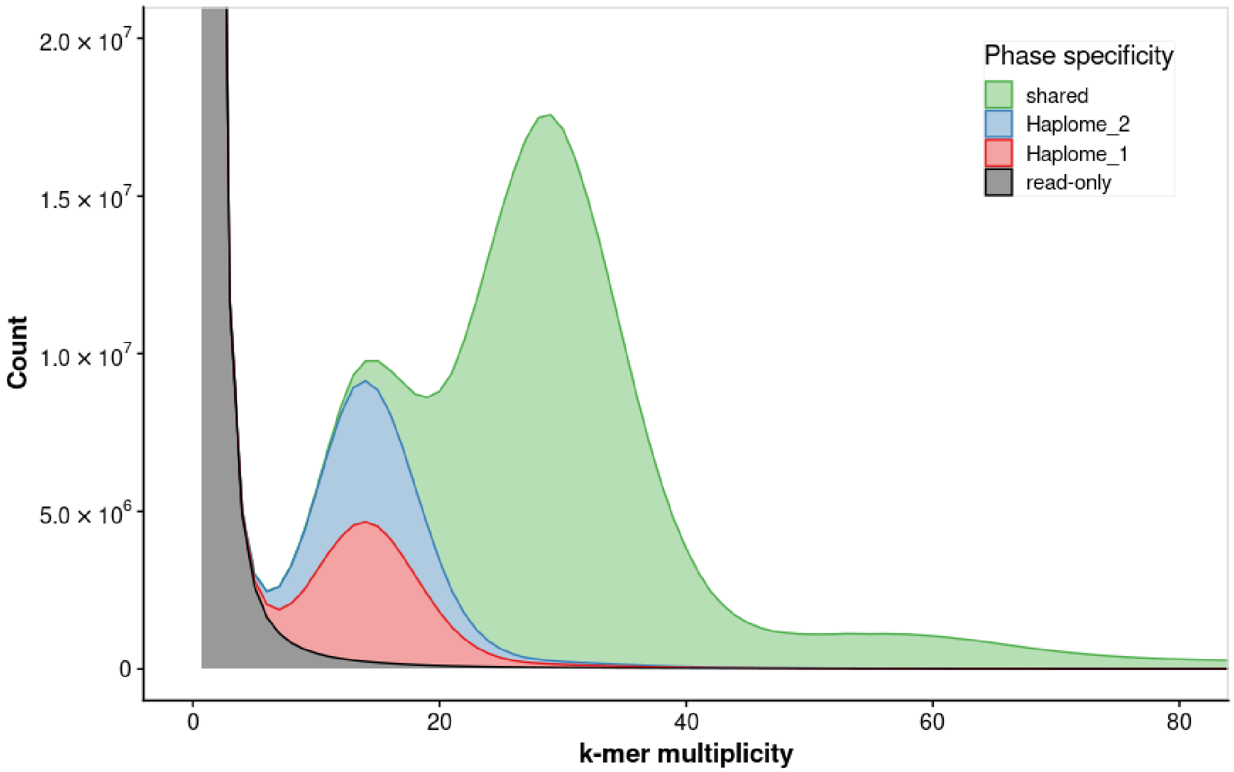


Supplemental Figure 1. Merqury *k*-mer multiplicity analysis of the phased chromosome-only assembly.

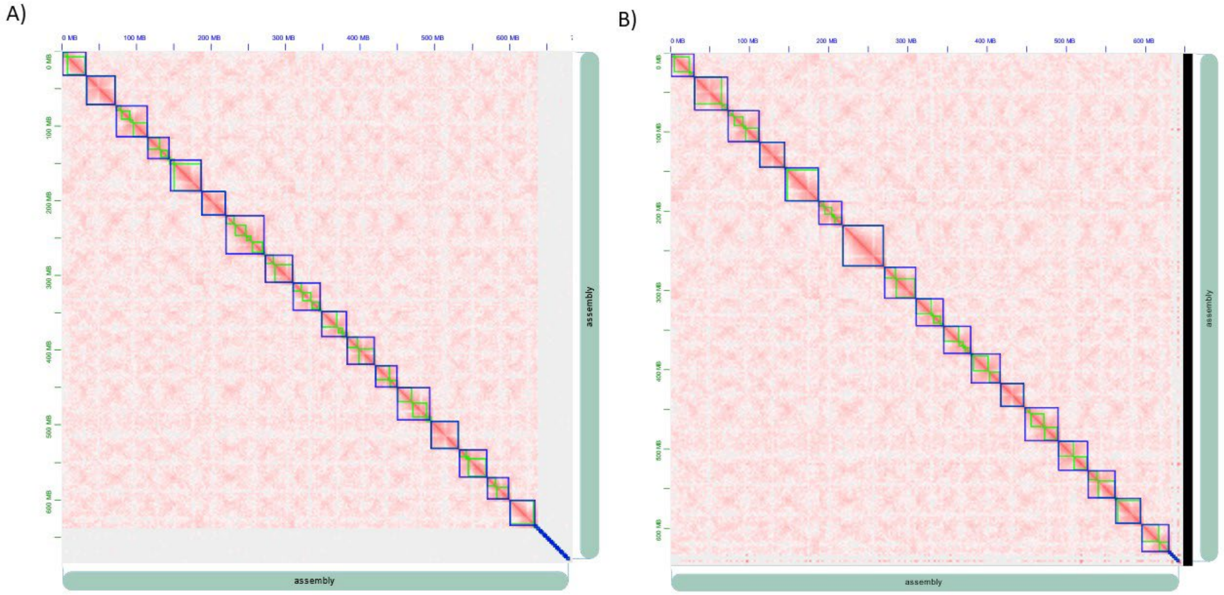


Supplemental Figure 2. Hi-C contact maps with the scaffolded genomes. A) Haplotype 1. B) Haplotype 2.

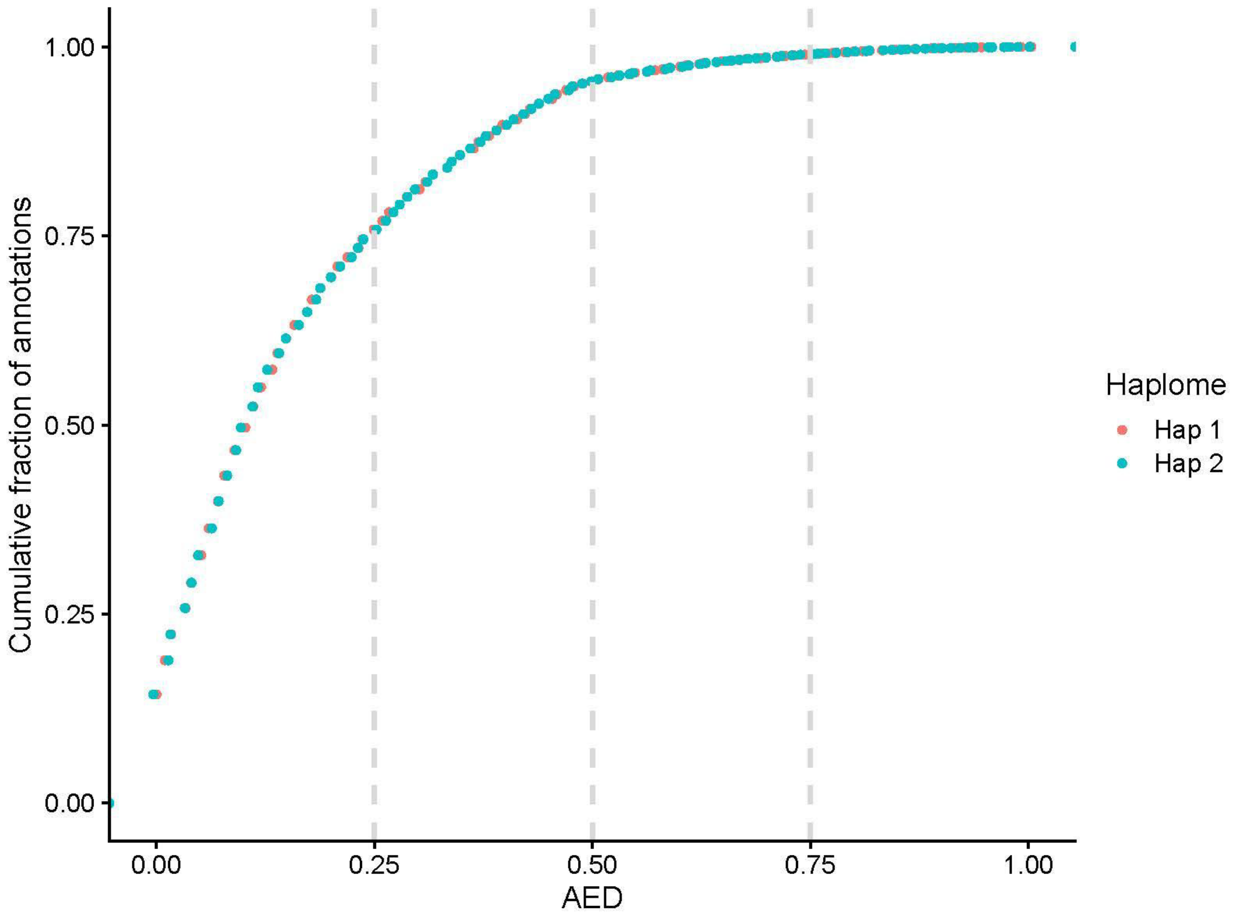


Supplemental Figure 3. Annotation Edit Distance (AED) of the gene annotation for Haplome 1 and 2.

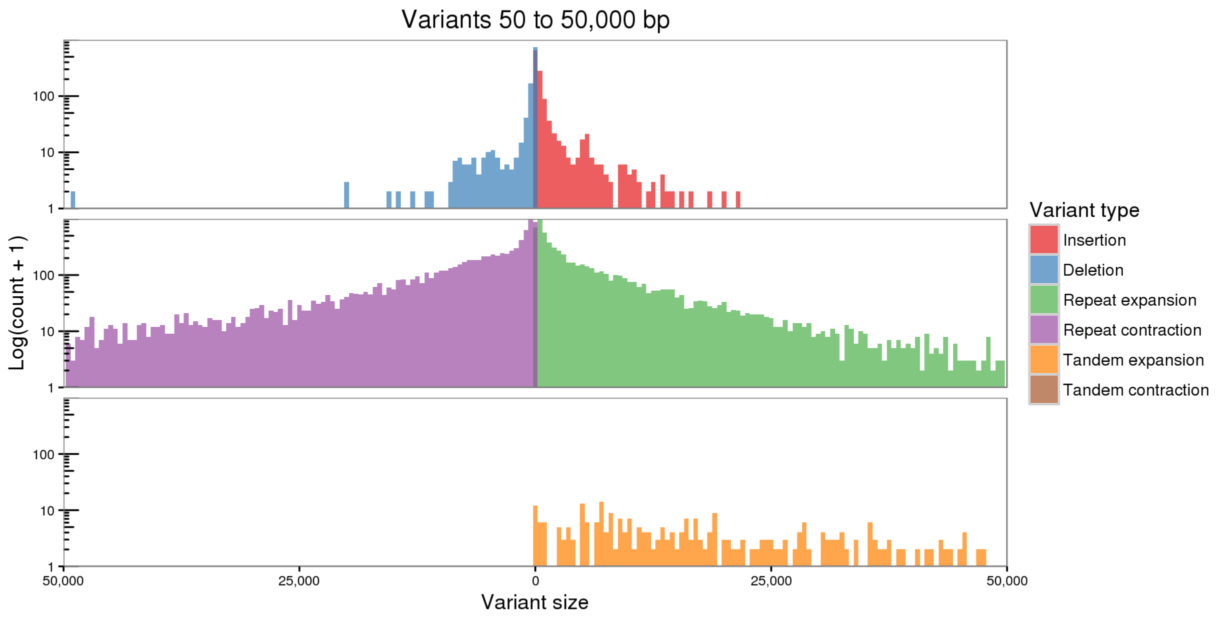


Supplemental Figure 4. Assemblytics structural variation between *M. fusca* and *M.* x *domestica* GDDH13.

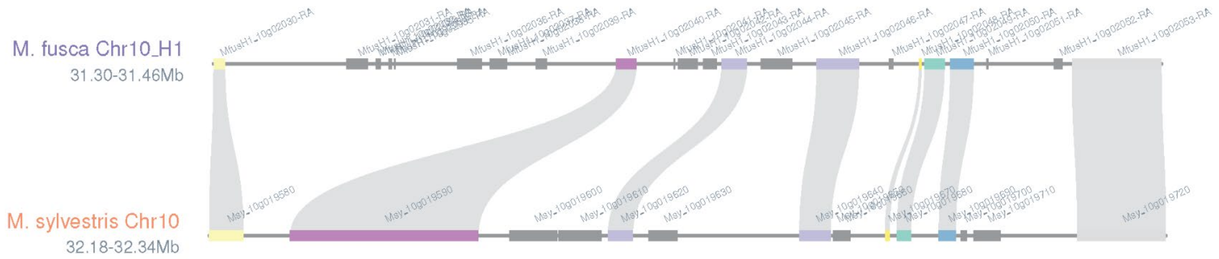


Supplemental Figure 5. Synteny within the *Mfu10* locus associated with fire blight resistance in *M. fusca* (top) compared to *M. sylvestris* (bottom). Orthologs of G-type lectin S-receptor-like serine/threonine-protein kinases genes (R-genes) are annotated by locus id and syntenic genes are annotated by color between the four genomes.
